# Windows of opportunity for Ebola virus infection treatment and vaccination

**DOI:** 10.1101/125336

**Authors:** Van Kinh Nguyen, Esteban A. Hernandez-Vargas

## Abstract

Ebola virus (EBOV) infection causes a high death toll, killing a high proportion of EBOV infected patients within 7 days. Comprehensive data on EBOV infection are very fragmented, hampering efforts in developing therapeutics and vaccines against EBOV. Under this circumstance, mathematical models become valuable resources to explore potential controlling strategies. In this paper, we employed experimental data of EBOV-infected nonhuman primates (NHPs) to construct a mathematical framework for determining windows of opportunity for treatment and vaccination. Considering a prophylactic vaccine based on recombinant vesicular stomatitis virus expressing the EBOV glycoprotein (VSV-EBOV), we found that the time window can be subject-specific, but vaccination could be protective if a subject is vaccinated during a period from one week to four months before infection. For the case of a therapeutic vaccine based on monoclonal antibodies (mAbs), a single dose might resolve the invasive EBOV replication even it was administrated as late as four days after infection. Our mathematical models can be used as building blocks for developing therapeutic and vaccine modalities as well as for evaluating public health intervention strategies in outbreaks. Future laboratory experiments will help to validate and refine the estimates of the windows of opportunity proposed here.

## Introduction

Emerged in 1976, Ebola virus (EBOV) has since caused numerous outbreaks in West African countries infecting ten to hundreds of cases^1^. The recent outbreak in West-Africa (2014-2016) resulted in nearly 30.000 infected cases with one-third of them being fatal^2^. Damages to the vascular systems during infection lead to bleeding, multiorgan failure, hypotensive shock, and death^3^. Ebola virus disease (EVD) displays in a wide range of non-specific symptoms early after infection, making diagnosis and early detection difficult^3^. The infection is acute leading to death within one to two weeks^1,3^. As a result, complete observations of disease progression or comprehensive evaluations of potential treatment options are problematic.

Experimental observations showed that the immune system often fails to control EBOV infection leading to elevated levels of viral replication^3^. Adaptive immune responses were poor or absent in fatal cases while survivors developed sustained antibody titers^3^. However, follow-up durations were different between fatal cases (approximately one week^1^) and survivors (from a few weeks to months^1^). Currently, treatment of EBOV infection is mainly based on supportive care^4^. Vaccine and therapeutics approaches are still under development and licensure with promising results for certain antivirals^4–6^, passive immunotherapy and vaccinations^7,8^. On one hand, EBOV infected nonhuman primates (NHPs) treated early with monoclonal antibodies (mAbs) were able to recover after challenged with a lethal dose of EBOV^9^. Furthermore, EBOV infected human treated with mAbs in addition to intensive supportive care were more likely to recover^4^. On the other hand, macaques vaccinated early with the VSV-EBOV vaccine survived lethal EBOV challenge^10^. Based on VSV-EBOV vaccine, a recent community trial showed protective efficacy in a ring vaccination approach^11^. These results prompted that the outcome of EBOV infection is sensitive to the time of intervention. Failing to catch up with the infection course could alter the chance to survive EBOV infection. Tailoring time windows of intervention is thus critical at both clinical and epidemiological levels.

Building a tractable approach that integrates systematically biological and medical research data is crucial to harness knowledge and to tailor therapies and vaccines. In this context, mathematical modeling has been a useful companion approach to advance understandings on mechanisms behind incomplete empirical observations. An overwhelming amount of modeling studies have been done in influenza virus^12–17^, human papilloma virus (HPV)^18^, and human immunodeficiency virus (HIV)^19–21^. These studies provided interpretations and quantitative understandings of the mechanisms that control viral kinetics, which are instrumental to formulate treatment recommendations^21–25^. Although West-Africa countries have been agitated by EBOV infection for decades, modeling studies of EBOV infection are rare. To the best of our knowledge, the first endeavor to model EBOV replication in *vitro* showed that the EBOV’s basic reproductive number is at least two fold higher than that for influenza virus^26^.

Using *in vivo* experiments in NHPs, this paper aims to model the interactions between EBOV replication and IgG antibody dynamics with and without passive immunotherapy. In particular, variations of simple mathematical models representing different interaction mechanisms were fitted to selective parts of the experimental datasets. Goodness of fit of the models were compared when needed to rule out less supportive models. Developed models were then used to estimate the needed time windows to achieve effective interventions. Considering an EBOV infection with a high infective dose just after vaccination, our numerical results showed that regular antibody response dynamic by vaccination would rather be late to control EBOV replication. To prevent a lethal infection outcome (i.e. viral load higher than 10^6^ TCID^50^), a host needs either a high antibody concentration early after infection or an alternative therapy in-place sufficiently early to enable the host’s adaptive immune responses to catch up the infection. Simulations of the developed models provided estimates for these critical windows of opportunity. In particular, therapeutic treatment could be effective if an assumed long-lived monoclonal antibody was administrated up to day 4 post infection. Prophylactic vaccination can be protective if it was given at least 6 days before exposure. However, circulating EBOV-specific antibody could diminish below protection level approximately three months after vaccination. Altogether, the framework presented here could help to tailor appropriate time windows for effective therapeutic and prophylactic interventions.

## Results

### Study Design

To construct mathematical models for determining windows of opportunity for both treatment and vaccination, we considered data from the two complementary strategies: a passive^9^ and an active immunization intervention protocol^10^. Schematic representation of the NHP experiments are provided in Fig. 1. Additional experimental details can be found in Materials and Methods. Viral load data in the controlled and treated cases were extracted from both the studies^9,10^. Antibody dynamics (IgG) data were available to assess its effects on the viral load^10^ whereas effects of mAbs were only available in terms of administrated time points and dosages.

Note that NHPs data is considered the best animal model to recapitulate EVD observed in humans^27^. In addition, controlled and defined experimental conditions are key guidances to model the complex interactions between virus and immune responses, e.g., defined time of infection and innoculum, consistent sampling time among the subjects, uniform host’s conditions, and consequently immune responses. The viral load was considered to determine the effect of intervention strategies. Epidemiological and pharmacological studies reported that a viral load higher than 10^6^ copies/mL^4,28^ is associated to a higher mortality rate, whereas observations on experimental data in NHPs showed animals with viral load levels higher than 10^6^ TCID_50_ were fatal^9,10^. Thus, we assumed that subjects with viral load levels higher than this threshold will have a *severe outcome*.

**Figure 1.**
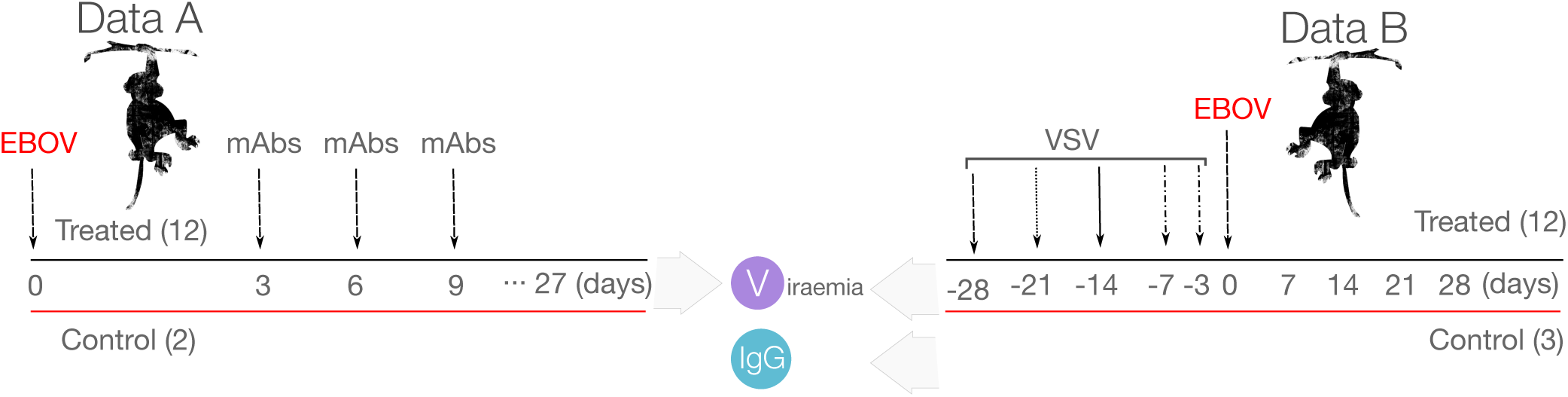
Experimental designs of the two data sources. The experiments were conducted in non-human primates. Number in the brackets is the sample size; *mAbs:* time of monoclonal antibody treatments^9^; *VSV*: time of vaccinations with vesicular stomatitis virus (VSV)^10^; *EBOV*: time of Ebola virus infection.

### Antibody profile after EBOV vaccination

To avoid ending up with a complex interaction model, IgG data were modeled independently to have a general profile of IgG responses, which is used later as an input in models of viral replication. This was done by using the IgG dynamics data after EBOV vaccination but before EBOV infection challenge^10^. A typical immunogen dynamic can be summarized in two phases: a catabolic decay phase during which the antigen is taken up by macrophages and other phagocytic cells, and an immune elimination phase during which newly synthesized antibodies combine with the antigen forming antigen-antibody (AgAb) complexes which are also phagocytosed. These dynamics can be written as follows

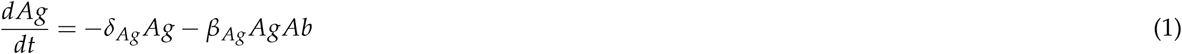

where *δ_Ag_* denotes the antigen removing rate. The parameter *β_Ag_* denotes the rate of AgAb complexes forming. The processed immunogen is then delivered to lymph nodes as the stimulus sources of the B-cell activation. This process can be written as an auxiliary delay state

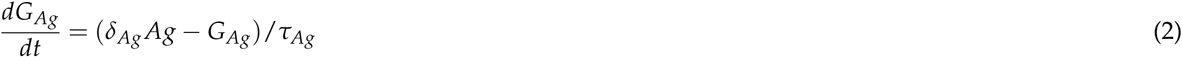

in which the processed immunogen is converted to the signal *G_Ag_* in time *τ_Ag_*. The above two processes lead to B-cell activation, proliferation, and antibodies secretion which can be summarized as follows

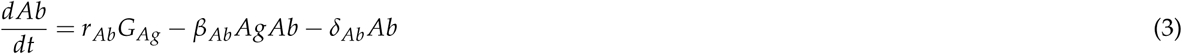

where *r_Ab_* reflects the end result of the three processes. The parameter *δ_Ab_* is the decaying rate of IgG which is approximately 28 days^29^. The parameter *β_Ab_* denotes the rate of AgAb complex forming. An ineffective antigen will not able to induce B-cell activation, i.e. *r_Ab_* = 0. Applying this to the IgG kinetic data^10^ results in a classical antibody response picture, namely a lag phase follows by an exponential phase before reaching a plateau (Fig. 2A). A high and steady level of IgG can only be acquired after two weeks. As a result, antibody responses may offer negligible protection during the first week after vaccination.

### EBOV replication profile

EBOV replication dynamics in the absence of any interventions were also modeled separately. This was done using EBOV titers (in TCID_50_) of only the control cases in the used datasets^9,10^. We considered two models, including the logistic growth model and a modified logistic growth model as follows

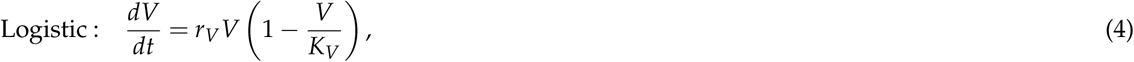

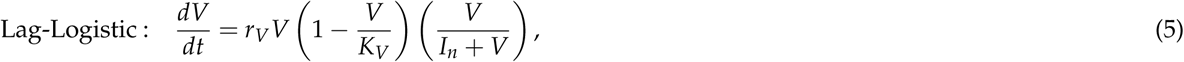

where *r_V_* denotes the virus replication rate and *K_V_* denotes the carrying capacity of the host. The parameter *I_n_* expresses a threshold below which the virus replication is restrained. Both models assume the viral replication is only limited by the available resources of the host. Considering a model selection based on AIC (see Materials and Methods), the model Eq. (5) with a lag-phase early after infection and slow growing phase (AIC=−10) portrayed the data better than the logistic growth model (AIC=21) in Eq. (4) (see also Figure 2B and the parameter estimates in Table S3). EBOV needed approximately three days to gain the momentum before growing exponentially, suggesting there is a crucial period for a successful treatment.

**Figure 2.**
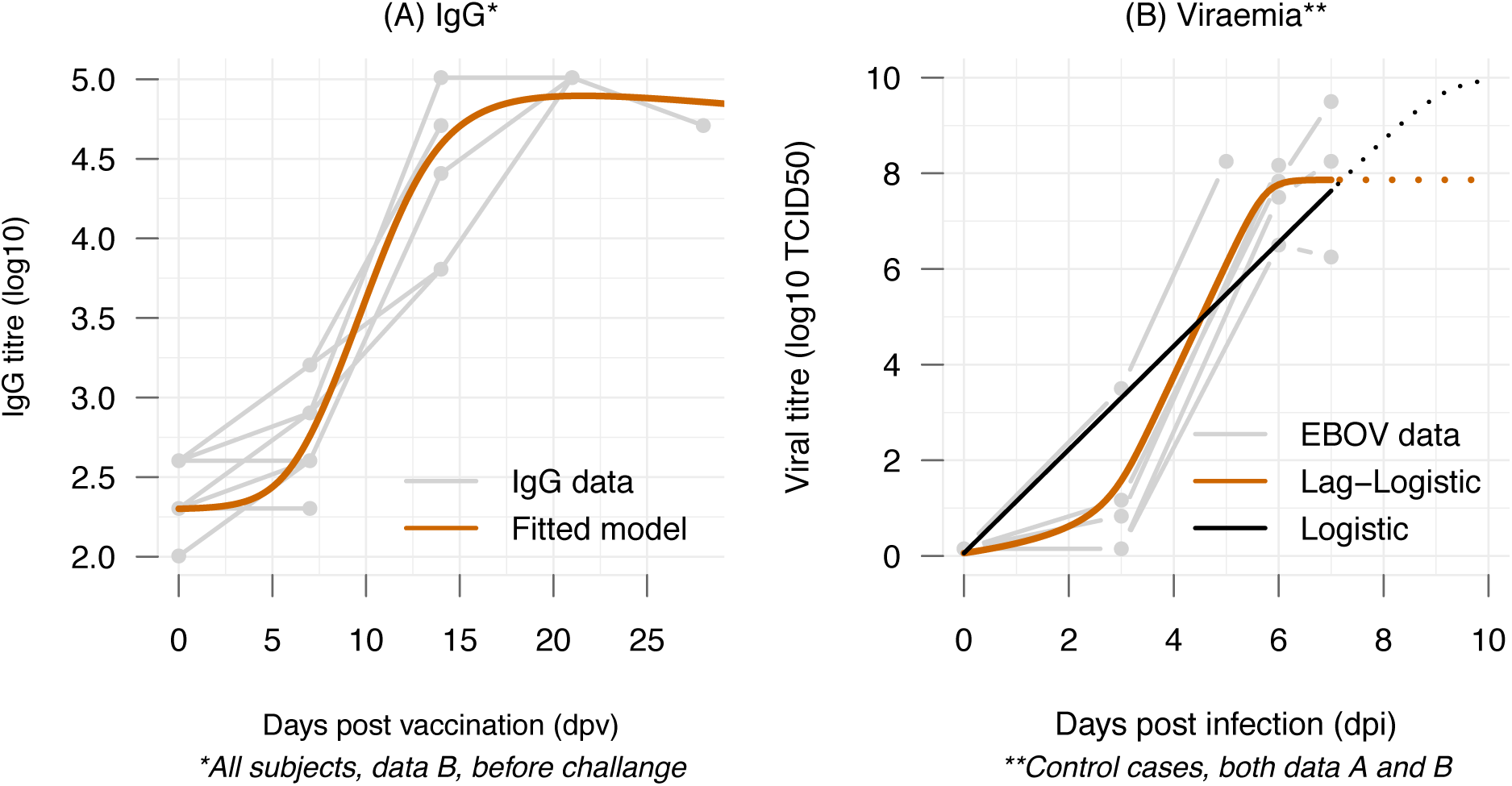
Fitting models to IgG and viral titer data. Gray lines are subject-specific data. **(A)** Data of IgG titer post vaccination and prior to EBOV challenge were used to fit to models of antibody responses. The fitted values of the models were superimposed (orange line) illustrating an average profile of IgG dynamics after exposure to EBOV. **(B)** Viral titers in control cases form both data sources^9,10^ were used to evaluate EBOV replication models in treatment-free scenarios. The follow-up data were stopped when the animals reached the endpoints to be euthanized^9,10^.

This result, in agreement with experimental observations, shows that even if the host develops a normal antibody response, EBOV would replicate unrestrained by the antibodies during the first week. As innate immune responses and consequently cellular adaptive responses were highly disrupted by EBOV^3^, this points towards the central role of antibodies in survivors of EBOV infection. Noting that the EBOV replication profile represents the cases infecting with a lethal dose, as such a varied, subject-specific lag-phase as a function of the innoculum can be expected. Nevertheless, in terms of safety, using EBOV dynamics based on a lethal dose to evaluate treatment therapies will provide the most conservative predictions.

### Tailoring windows of opportunity for prophylactic vaccines (VSV-EBOV)

To account for the effect of antibodies in controlling the virus, the viral replication model Eq. (5) was modified to

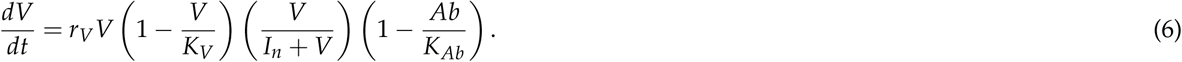

To connect to EBOV dynamics in (5), the parameter *K_Ab_* was introduced reflecting a functional threshold at which the antibody titers inhibit EBOV net growth rate. Crossing this threshold leads to the virus titer being cleared. To evaluate this model, IgG titer and viral load data of two animals vaccinated three days before EBOV infection were used (M31 and M32); these were the only two animals with detectable viral titers in the experiment^10^. Firstly, the antibody dynamics *Ab* were obtained by fitting the equations Eq. (1)–(3) to the IgG data of the two subjects. The parameters *δ_Ab_, δ_Ag_, β_Ag_*, and *β_Ab_* were fixed to the estimates derived earlier from the data of all subjects (Fig. 2A). The two parameters *τ_Ag_* and *r_Ab_* were refitted to allow subject-specific responses, see Table S2. Afterwards, the Ab outputs from the first step were used as inputs to fit the model Eq. (6) to viral titers data of the two subjects. With the assumption that EBOV would replicate indifferently among infected subjects, the parameters *K_V_* and *I_n_* were fixed to the previous estimates from the model Eq. (5).

Figure 3A-C shows that the models Eqs. (1) to (3) rendered faithfully the IgG dynamics in the three animals. Differences observed in the dynamics can be explained by subject-specific responding time to stimulate B cells (*τ_Ag_*) and to produce EBOV specific antibodies (*r_Ab_*), see Table S2. Figure 3D-E show that the model Eq. (6) reproduced the viral dynamics in the two subjects (M31 and M32). The differences in IgG dynamics also lead to different working threshold estimates of *K_Ab_* for each subject, reflecting possibly different antibody responses strength.

The model of the interactions between IgG and EBOV allow to simulate windows of opportunity for vaccination. By varying the time of vaccination, a time period during which a vaccine administration could prevent a likely-lethal viral load level can be estimated. Since the threshold for a functional antibody response (*K_Ab_*) can be subject-specific (Fig. 3F), a range of thresholds based on the observed IgG data from 10^2.5^ to 10^4.5^ were tested. Based on data of the control cases (Fig. 2B) and empirical observations in EBOV-infected human^28^, a subject expresses viral load level higher than 10^6^ could be considered as having a severe outcome. Figure 4 illustrates the time windows for different vaccination time and different working threshold *K_Ab_*. Noting while the chosen level can be subjective, one can simply lower the values to have a more conservative time window estimate.

It can be observed that the higher the working thresholds of antibody the shorter the time windows of intervention. For each *K_Ab_* threshold, there is a safe time window where viral titers were not observed. For example, a threshold *K_Ab_* = 10^4^ could prevent EBOV replication from reaching severe viral load levels if only the subject had been vaccinated at least 6 days before infection. However, if the subject had received vaccination for more than four months before, circulating antibody levels could have decreased below the working threshold (*K_Ab_*) at the time of infection. As such, if the secondary antibody responses to EBOV infection are not considerably faster than primary responses, the subject would also succumb to the disease. Here, the secondary antibody response to EBOV was assumed similar to a primary response, i.e. similar to the dynamic observed in Fig. 2A, and that the IgG titer were accumulative to primary response.

Remarkably, assuming an infected subject would develop the same IgG profile as a vaccinated subject, simulation results showed that a normal IgG response will fail to keep the viral load from reaching its peak, regardless of the working threshold *K_Ab_* (Fig. S1). At best, a normal IgG profile developing from the day of infection could clear the virus 9 days post infection, if the subject is still alive after several days withstanding massive viral titers.

**Figure 3.**
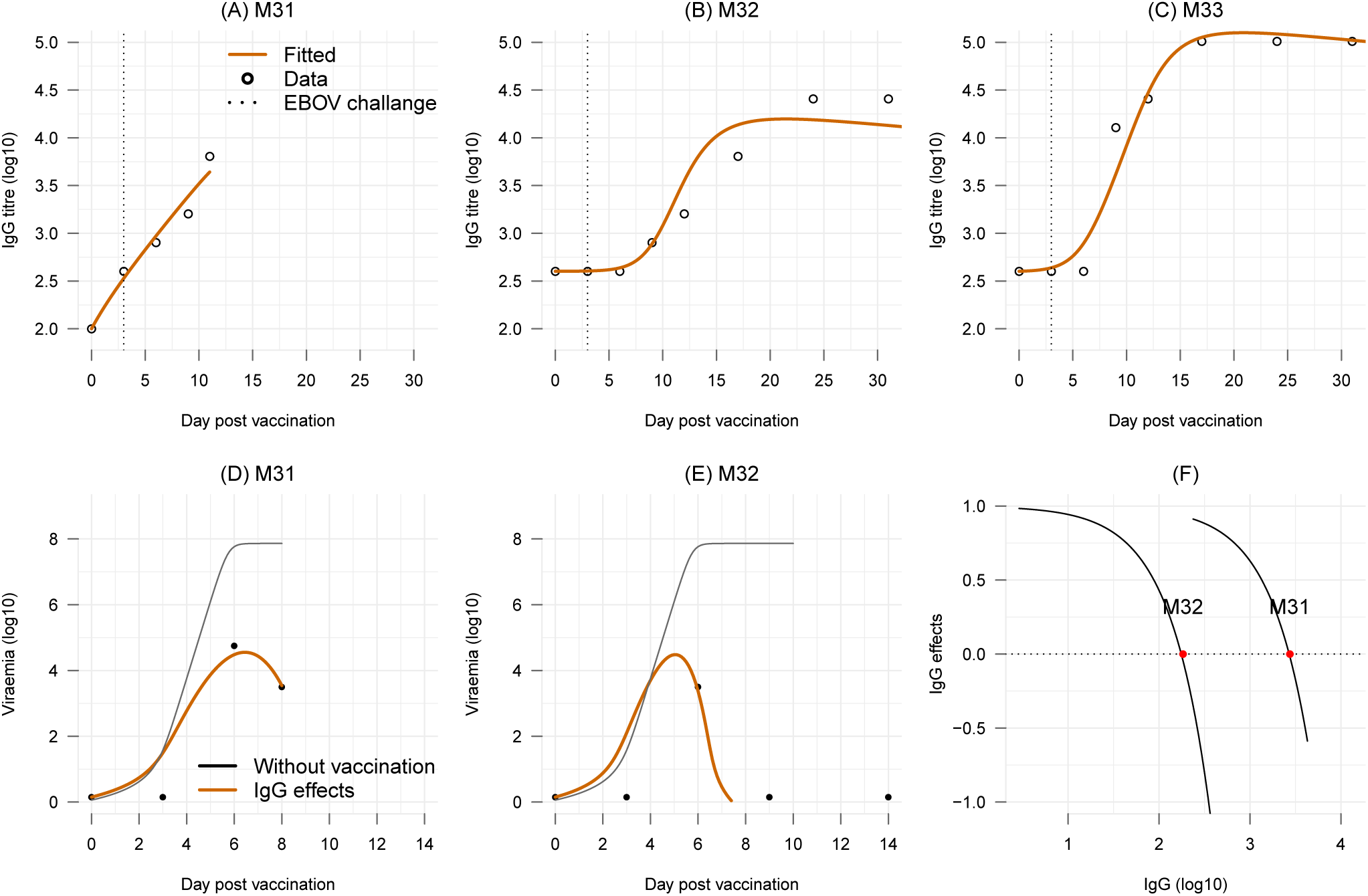
Effects of IgG antibody on controlling viral load. (**A-C**) Fitting IgG dynamics model Eqs. (1) to (3) to IgG data of the three subjects vaccinated three days before EBOV challenge. **(D-E)** Fitting viral dynamics model Eq. (6) to viral load data of the two subjects vaccinated three days before EBOV challenge. The model without vaccination (solid black line) is added as reference. **(F)** Functions of IgG effect on controlling viral growth in each subject.

### Tailoring window of opportunity for therapeutic vaccines (mAbs)

In the experiment with passive antibody treatment^9^, viral dynamics in the animals treated with mAbs ceased after the first dose on day 3 post infection. Although this may include the role of host’s immune responses, the previous section has shown that a normal IgG profile starting at the day of infection may not not able to clear the virus at least until day 9 post infection (Fig. S1) and that antibody level were negligible during the first few days. Thus, for EBOV-infected subjects, the mAbs treatment would play a decisive role in tackling EBOV infection during the first days after infection. To recapitulate the viral dynamics in subjects treated with mAbs when antibody responses were negligible, the viral dynamics are rewritten as follows

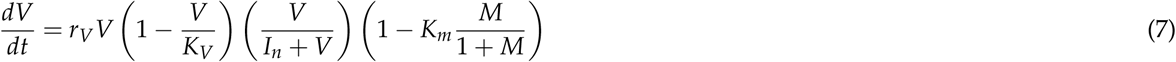

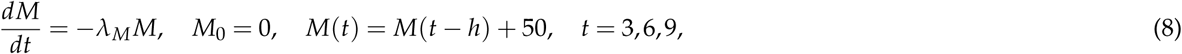

where *M* is the administrated dose of mAbs (see Materials and Methods) and *h* is the ODE integration step size. Here the mAbs are assumed acting indifferently from IgG antibody, i.e. not only reducing the viral replication but also promoting the viral clearance. The Eq. (3) expressed the assumptions that the mAbs concentration was accumulative over doses and decayed exponentially during the infection course. As a result, the combination of parameter *K_m_* and the dynamics of the mAbs leads to similar working threshold mechanism as in Eq. (6). Here, the parameter *K_m_* represents the maximum effect of the mAbs.

**Figure 4.**
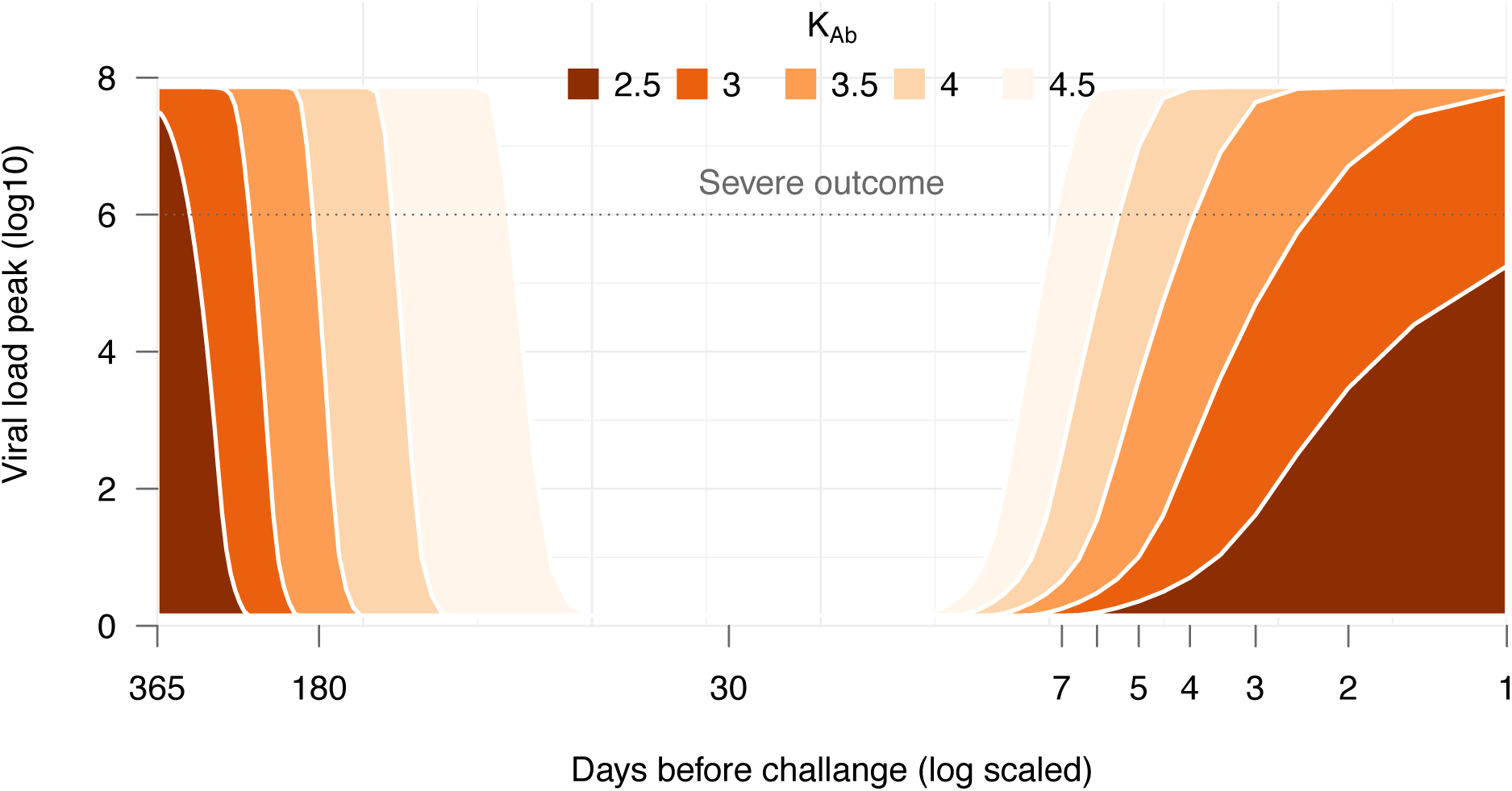
Simulation the effect of vaccination time and immune strengths. The time of vaccination before challenging a subject with a lethal dose of EBOV were varied from one year to one day. Secondary response of IgG dynamic was assumed similar and accumulative to the primary response. Colors represent the varied working threshold (*K_Ab_*). The smaller the threshold the stronger the effect of antibody in negating the viral replication. The model of viral dynamics in the presence of antibody (Eq. (6)) were simulated. The maximum viral load generated by the model in each combination of the vaccination time and the working threshold was retrieved and plotted.

Evaluation of the model Eqs. (7) to (8) were done by fitting to the viral dynamics data, separately for each treated animal with observable viral load^9^. As the model is ignoring the effects of antibody responses, only viral load data at time points from day 0 to day 9 were used. The parameters *r_V_, K*_*V*_, and *I_n_* were fixed to the earlier estimates in Eq. (5). For simplicity, the mAbs are assumed to have a stable and long elimination half-life of 28 days across the subjects^30^.

Figure 5 shows that the model portrays adequately the viral load kinetics in every subjects. Interestingly, the 134 mAbs treatment effect seemed to be separated in two groups: a low effect group (*K_m_* < 1) that allowed viral titers to 135 linger until day 9 and a high effect group (*K_m_* > 1) that quickly stemmed down the viral titers (details in Table S3). Extrapolating the model Eq. (7) to time points after day 9 post infection showed a sustained viral load in those subjects whose mAbs effect is low (*K_m_* < 1), see Fig. 5. Because the mAbs was already assumed having the longest elimination half-life observed in natural antibody, this result suggests that mAbs treatment alone may be insufficient for those the mAbs effect is low.

**Figure 5.**
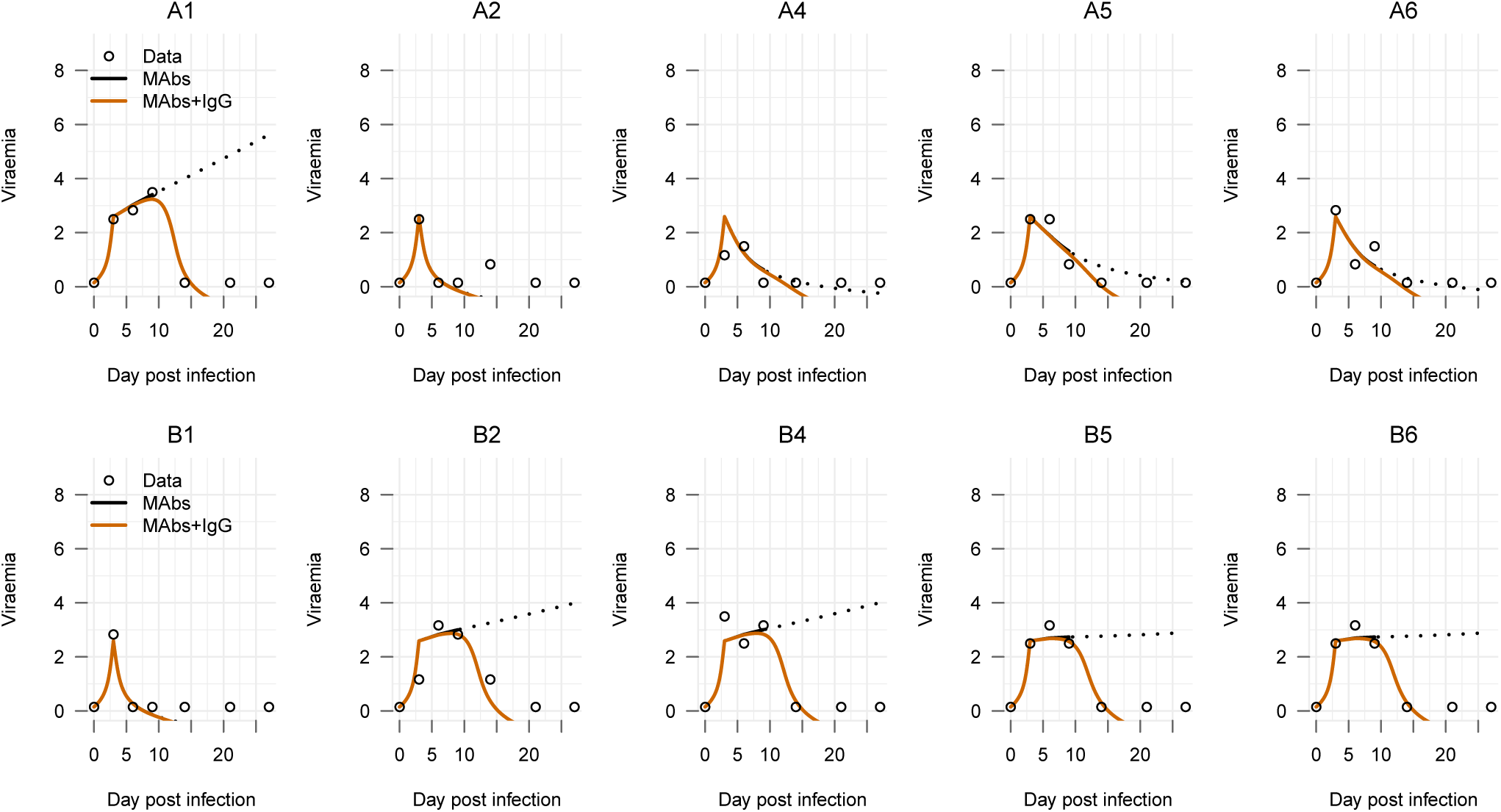
Fitting the mAbs treatment effect model. *mAbs:* fitted model with only mAbs effect during the first nine dpi, dashed line shows the extrapolated viral load kinetics from this model; *mAbs-IgG:* adding the general IgG profile with the working threshold *K_Ab_* = 10^4.5^. mAbs half-life is 28 days. Two different combinations of monoclonal antibodies were tested in NHPs (ZMapp1 and ZMapp2)^9^. The top panel of figures (A1 to A6) presents the six NHPs receiving three doses of ZMapp1, while the bottom panel of figures (B1 to B6) presents the six NHPs receiving three doses of ZMapp2^9^.

To take into account the effect of host’s IgG response, we incorporated the general IgG profile developed earlier into the model Eq. (7) and simulated the viral load dynamics with a conservative working threshold 10^4.5^ in each subject as follows

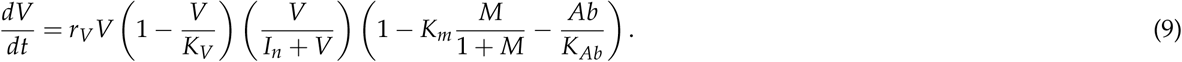

Here it was continued to assume that EBOV-infected subjects would able to develop a similar IgG profile as in those 141 subjects vaccinated with VSV-EBOV^10^. Figure 5 shows that including the effects of IgG into the treatment model 142 replicated closely the viral load data. Therefore, the host’s antibody response (IgG) would have played the key role in resolving the infection for those mAbs treatment were not sufficient.

In light of these results, therapeutic treatment windows can also be developed using the model Eq. (8) and Eq. (9). For illustration purpose, we varied the time of treatment administration to define which treatment initiation can prevent viral load to reach fatal levels, i.e., viral load is higher than 10^6^ TCID_50_. Figure 6 illustrates EBOV kinetics considering a single dose treatment approach, we can observe that a single dose of a long-lived mAbs administrated up to day 4 post infection was able to clear the virus before it reached the likely-lethal viral load.

**Figure 6.**
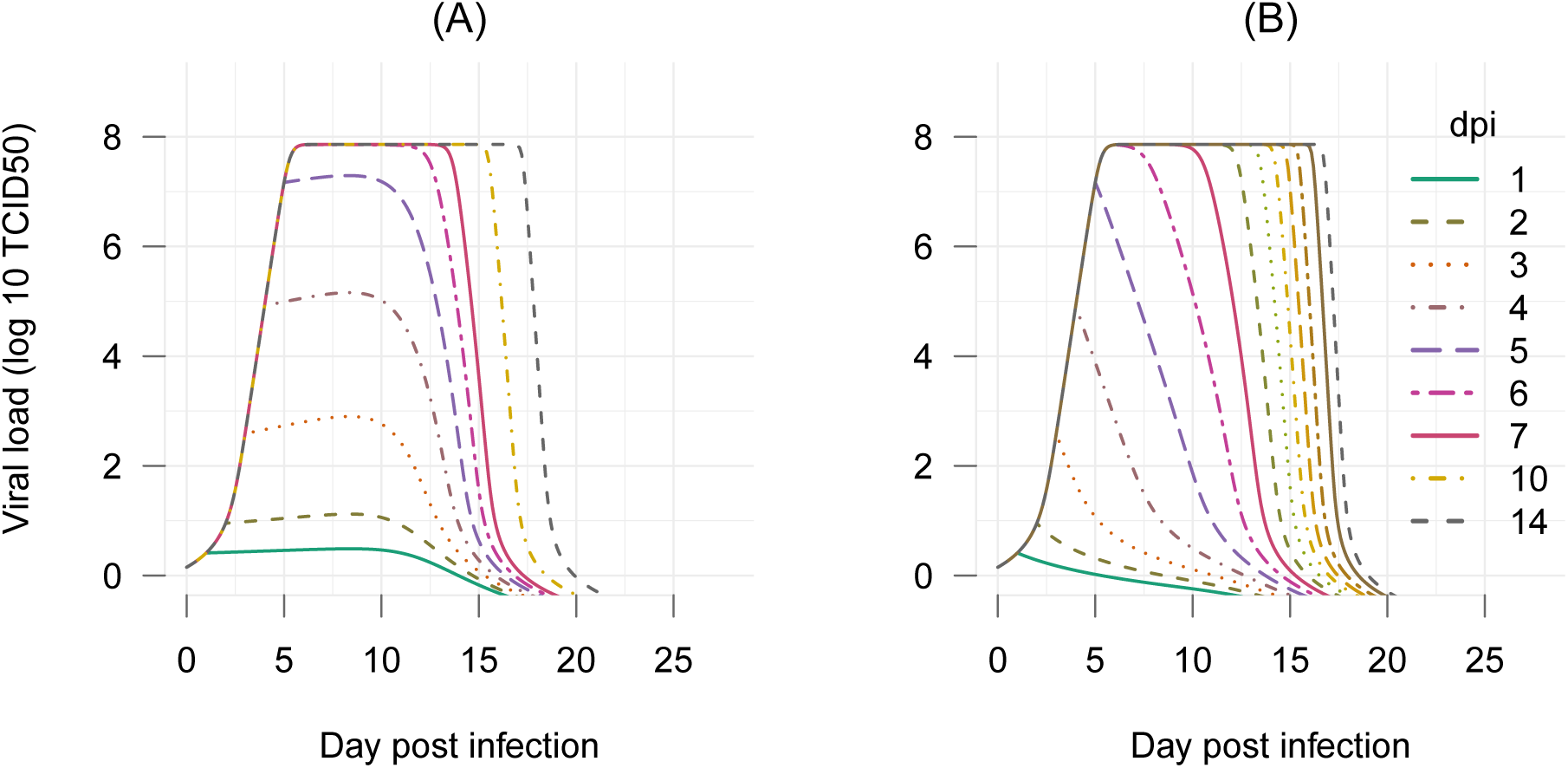
Simulation of single-dose mAbs treatment assuming a long mAbs half-life of 28 days. The time administrated mAbs were varied from 1 to 14 days. For each regimen, the model of viral dynamic including both treatment and IgG effects were simulated to generate the corresponding viral load dynamics. (A) assuming low effect (*K_m_* = 0.98), normal IgG response profile with working threshold *K_Ab_* = 10^4.5^, and long half-life mAbs of 28 days; (B) assuming high effect (*K_m_* = 1.47), normal IgG response profile with working threshold *K_Ab_* = 10^4.5^.

## Discussion

The recent unprecedented EBOV outbreak in West Africa has affected more people than in all previously outbreaks combined. While significant progress has been made in therapeutics and vaccines against EBOV on preclinical level, no licensed products are currently available. Lack of market for products, consequently interests of pharmaceuticals companies could have hindered the progress. Furthermore, the high pathogenicity of EBOV hampers the possibilities to have comprehensive data and to conduct clinical and efficacy studies in EBOV infection. In this context, using mathematical models in combination with experimental data can be essential for EBOV countermeasure development.

Our results suggested that even if antibodies would response normally to vaccination at day of infection (Fig. 2A), the pace of antibodies response would still be too slow to counteract EBOV replication, which needs only three days to start an exponential growth (Fig. 2B). In fact, antibody titers were negligible during the first week post vaccination. Therefore, protection to EBOV infection depends on having a high level of antibodies as all animals vaccinated sufficiently early were survived^10^. Future experiments should closely monitor IgG dynamics in infected versus vaccinated cases. Comparing the responding time in the two situations will help to clarify if the adaptive immune responses are indeed malfunction.

Differences in the race between the EBOV replication and the immune system response highlight the importance of timeliness in EBOV treatment. By variations in the time of vaccine administration, simulations showed that the window of opportunity for an effective intervention is limited. Beside the pathogen replication dynamic, key parameters to the window estimates are the time to vaccination responses, the pathogen-specific antibody half-life, and secondary antibody responses to the pathogen. As of now, lack of data about EBOV reinfection does not allow to obtain more accurate estimates. Future controlled experiments in NHPs can evaluate the memory of immune responses to EBOV reinfection to enable detailed evaluations of EBOV vaccination strategies. It is worth noting that our models have not taken into account the effect of prime-boost immunization protocols which could significantly widen the left boundary of the time windows. Prime-boost strategies have yielded 30-fold or greater increases in antibody titers^31^.

Combination of mAbs represents one of the most promising therapeutic modality^4^. Our results showed that early use of this supportive treatment is crucial in preventing a fatal outcome. However, subject-specific responses to the mAbs can be expected. When the role of a host’s antibody response were neglected, mAbs treatment can clear the virus in some but not all the subjects (Fig. 5). Both, a long and a short half-lives, exhibited the possibility of a viral rebound if the antibody host responses were neglected (Figs. 5 and S2). These results reiterate the key role of the host’s antibody response in clearing the virus once the treatment effect wears off.

As the elimination half-life estimate of the used mAbs were not reported, it was not possible to narrow down the estimates of the drug effect, consequently the estimates of the time window. For example, a half-life of half an hour can also produce the viral load data (Fig. S2) but the estimated drug effect were rescaled (Table S3). However, this can be easily overcome when common pharmacokinetics parameters are available, such as the elimination half-life for the mAbs. This approach opens the opportunity to mathematically evaluate EBOV treatment regimens. Our example of assessing a one-dose regimen illustrates that highly beneficial information can be obtained.

Note that the windows of opportunity reported here based on data of NHPs infected with a lethal infective dose, thus it would represent the infection in fatal cases. Depending on the infective dose and individual responses strength (*K_Ab_*) in each situation, the windows will become wider or narrower. Varying these conditions and simulating the viral load showed that the validity of the models were supported when the generated viral load dynamics exhibits known characteristics of EBOV infection in human (Fig. 7). For example, the time from infection until the exponential growth of EBOV ranges from 2.6 to 12.4 days (median: 3.8), which is equivalent to the incubation period of EBOV in human^1^. Additionally, the time from from infection to death ranges from 8.1 to 15.1 days (median: 9), and the time to recovery ranges from 6.9 to 17.6 days (median: 9.7) also resemble to that observed in practice^1^. Based on these kinetics, infected subjects could develop the disease after several days having no detectable viral load. This suggests that EBOV treatments need to provide as early as possible for all those exposed, even that they express no signs or symptoms. Cares and quarantine procedures would also need to be provided for exposed individuals at least twenty days which is twice the maximum incubation period estimated above.

**Figure 7.**
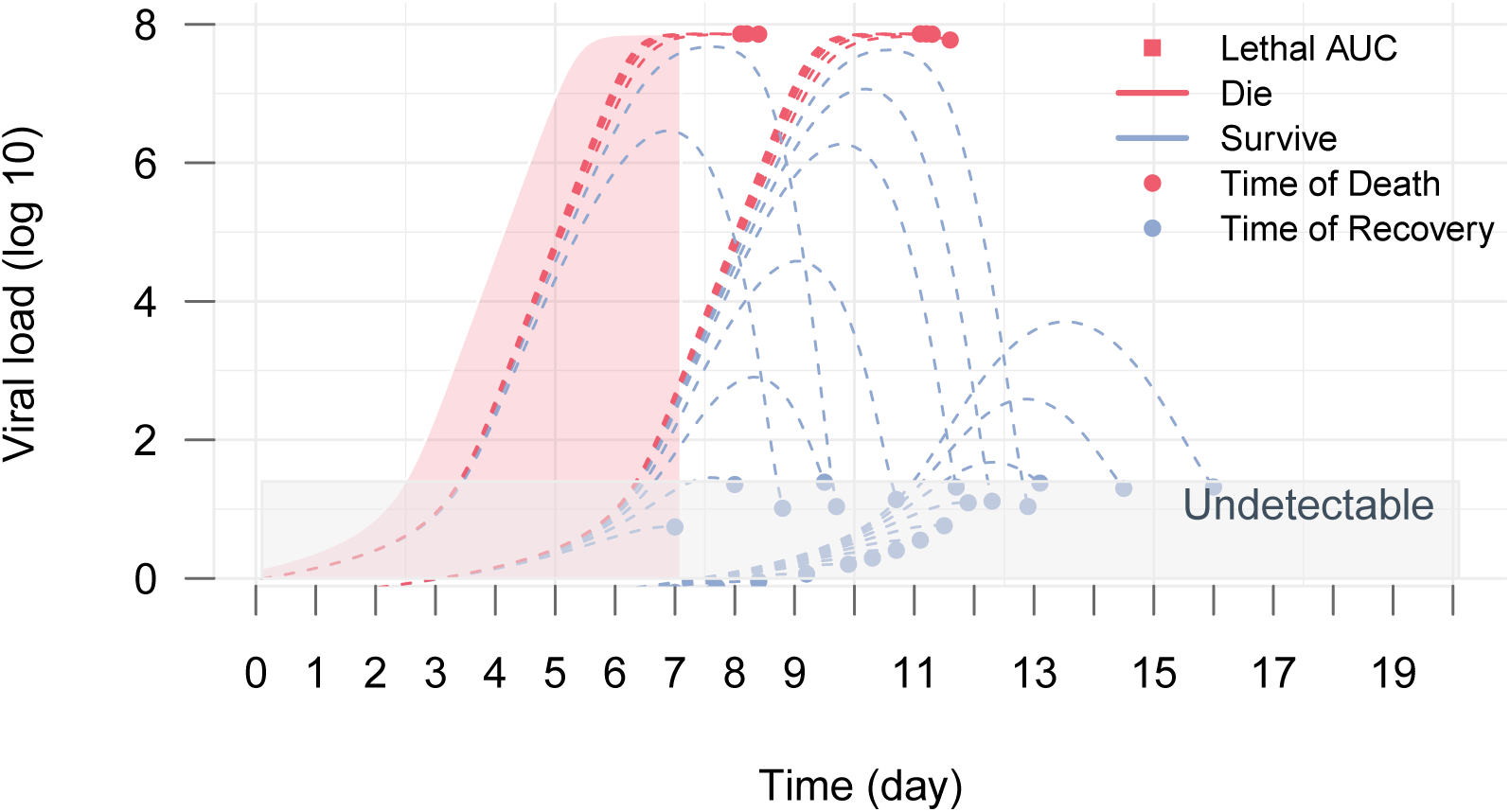
Simulations of the viral replication model for different infective doses and the immune responses strengths. The red area depicts the total viral load (AUC) that is used at the reference threshold for lethal criteria. This equals to the total viral load in control cases presented in Fig. 2. Subjects withstand a total viral load higher than the threshold is assumed fatal, otherwise they will recover once the viral load is resolved under detection level.

In summary, this paper proposed mathematical models by using selective parts of different data sources for model evaluation, resulting in a general framework for the development of treatment regimens and vaccination strategies. On top of that, public health policies and initiatives can also be evaluated with realistic treatment efficacy and subject-specific immune responses. In the scarcity of data, mathematically modeling approaches posits a strong potential to uncover useful information in controlling infectious diseases which is gradually become pivotal in the years to come.

## Materials and Methods

### Experimental data

Experimental data considering a therapeutic vaccine using monoclonal antibodies (mAbs) was taken from^9^. The mAbs were engineered to specifically recognize the EBOV glycoprotein (GP) inserted in the membrane of the viral particle. In this experiment, a group of 12 macaques were administrated mAbs intravenously at day 3, 6, 9 post infection with a constant dose of with 50mg/kg. These were divided into two groups, 6 NHPs received the monoclonal antibodies combination ZMapp1 (Group A) and the other 6 received ZMapp2 (Group B)^9^. No treatment was given to the two control cases. EBOV titer increased rapidly but ceased when the first dose of mAbs was administrated. The viral load continued to increase in the control cases until the subjects were euthanized at day 7 post infection. All animals cleared the virus from day 10 onward, with the exception of A1 which presented a high clinical score.

Experimental data for a prophylactic vaccine based on recombinant vesicular stomatitis virus expressing the EBOV GP (VSV-EBOV) was taken from^10^. In this experiment, groups of two or three macaques were vaccinated at 3, 7, 14, 21, and 28 days before EBOV challenge. Macaques were immunized with a single intramuscular injection of plaque-forming units (PFU) of VSV-EBOV. An ineffective vaccine (the VSV-Marburg virus vaccine (VSV-MARV)) was given to three control cases. IgG titers were measured regularly four weeks before and after the challenge. All the vaccinated animals showed a sharp increase of IgG titers one week after vaccination. IgG titers sustained at the level above up to two months. EBOV titers were monitored up to 9 days after the challenge. All the control cases showed a high level of viral titers and were euthanized five to seven days after infection. Viral titer was not observed in all the animals vaccinated at least seven days before the challenge. Among three animals vaccinated three days before the challenge, two had observable viral load in which one died and the other survived. A schematic representation of both NHPs experiments is provided in Fig. 1.

### Model fitting and selection

Selective parts of the datasets were used to evaluate models representing different mechanisms. When applicable, model comparison was done using Akaike information criteria (AIC). When data are available, extra components involved in the models were computed as forcing functions by linear approximation. These functions were used as inputs in model fitting instead of adding extra model equations. Time points when there were no measurable viral load were imputed as 10^0.15^ TCID_50_^32^. Model fitting was conducted in log ten for both the states and parameters. Objective function was defined as the root mean square error of the fitted value and the experimental data. Optimization was done with the Differential Evolution algorithm using the recommended configurations^33^. All simulations were done using R^34^. Details of model fitting can be found in Table S1.

## Acknowledgements

The authors thank Andrea Marzi and Heinz Feldmann (Laboratory of Virology, NIAID, NIH) for providing data from previous animal studies. This work was supported by the Alfons und Gertrud Kassel-Stiftung and iMed-the Helmholtz Initiative on Personalized Medicine. VKN has been supported by the President’s Initiative and Networking Funds of the Helmholtz Association of German Research Centres (HGF) under contract number VH-GS-202.

## Author contributions statement

VKN and EAHV designed the modeling. VKN performed the simulations. VKN, EAHV analyzed the data. VKN, EAHV discussed and wrote the manuscript. All authors reviewed the manuscript.

